# Alpha Oscillations Encode Bayesian Belief Updating Underlying Attentional Allocation in Dynamic Environments

**DOI:** 10.1101/2023.08.24.554581

**Authors:** Siying Li, Carol A. Seger, Meng Liu, Wenshan Dong, Wanting Liu, Qi Chen

## Abstract

In a dynamic environment, expectations of the future constantly change based on updated evidence and affect the dynamic allocation of attentional resources.To further investigate the neural mechanisms underlying efficient allocation of attention, we employed a modified Central Cue Posner paradigm in which the probability of cues being valid (that is, accurately indicated the upcoming target location) was manipulated. The proportion of attentional resources allocated to the cued location (α), which was sensitive to precision of predictions on previous trials, was estimated using a hierarchical Bayesian model and was included as a regressor in the analyses of electrophysiological (EEG) data. Our results revealed that before the target appeared, the attentional modulation of alpha oscillations (8∼13Hz) for high-certainty cues (88% valid) were significantly predicted by the attentional resources measure (α). This relationship was not observed under low-certainty conditions (69% and 50% valid cues). After the target appeared, the attentional resources measure (α) correlated with alpha band oscillations only in the valid cue condition and not in the invalid condition. These results provide new insights on how trial-by-trial Bayesian belief updating relates to alpha band encoding environmentally-sensitive allocation of visual spatial attention.

## Introduction

Anticipating upcoming events significantly impacts how we respond, especially when confronted with a volatile and uncertain environment (Boehm, 1994; Jongen and Smulders, 2007). Our decision-making and behavioral responses are influenced by our prior beliefs formed through past experiences, as well as our capacity to adapt flexibly to ever-changing circumstances (Blennow et al., 2020; Sambrook et al., 2021). Previous research has demonstrated that prior beliefs can influence sensitivity, processing, and performance in perceiving stimuli, underscoring the central role of expectations in shaping behavior (Adlard et al., 2008; Hein et al., 2023; Hein and Herrojo Ruiz, 2022; Kok et al., 2013). The variable possible states of a volatile environment are key factors in shaping prior beliefs. To better understand the role of environmental fluctuations in attention, several studies have employed computational models in conjunction with fMRI, EEG, and other neural recording techniques to demonstrate that the fluctuating environment is a critical factor that affects neural activity underlying attentional modulation (Hsu et al., 2005; Yu and Dayan, 2005).

In the Central Cue Posner paradigm, spatial cues are probabilistically related to target location. Over trials, participants form prior beliefs regarding the validity of the cue: that is the probability that the cue will correctly indicate the target location (Posner, 1980). Behavioral studies have revealed that increasing the proportion of valid cues leads to a larger difference in reaction time between valid and invalid trials (Risko and Stolz, 2010). Using fMRI techniques, numerous studies have identified specific responses in the posterior and temporal-parietal regions to valid cues (Corbetta et al., 2014, 2008). Vossel et al. (2006) found that attentional resources were deployed to the cued location and withdrawn from the uncued location with high validity cues, and these changes in attentional resource deployment were encoded by a right frontoparietal attention network. To summarize, the sensitivity of attention to probabilistic changes is reflected not only in behavioral performance but also in neural activity. In this study, we utilized a combination of computational modeling and electrophysiology (EEG) to examine how changes in the probability of validity for cues impacts the deployment of spatial attention, as encoded by alpha-band activity, and how these processes are influenced by prior beliefs.

Because the validity of the cues in the Central Cue Posner paradigm are initially unknown and can change over time, it is crucial to continuously infer the likelihood based on past experiences and observations. According to the Bayesian brain hypothesis (Knill and Pouget, 2004), the brain calculates the posterior probability by combining new information with prior beliefs, which are based on past experiences. This process of Bayesian belief updating allows the brain to make more accurate predictions and adapt its internal representations to the changing environment (de Lange et al., 2018; Zénon et al., 2019). The hierarchical Bayesian framework proposed by Mathys et al. (2011, 2014) has found extensive application in various research areas, including visual perception and learning (Kolossa et al., 2015; Körding and Wolpert, 2006; Liu et al., 2023; Yuille and Kersten, 2006). Multiple studies have demonstrated that this model consistently outperforms other simpler models, such as the Rescorla Wagner (RW) model (Liu et al., 2022; Vossel et al., 2015, 2014). Recently, there has been a growing interest in investigating the applicability of Bayesian frameworks in the encoding and decoding networks of attention (Sodkomkham et al., 2016; Visalli et al., 2021). However, in research pertaining to the spatial allocation of attentional resources, there is currently insufficient knowledge of the neural mechanisms underlying Bayesian inference and belief updating.

Alpha band activity is thought to be an effective neural correlate of underlying neural attentional mechanisms and can be used to track the allocation of attention across the visual field (Sutterer et al., 2021; Worden et al., 2000). Modern research has focused on exploring the inhibitory function of alpha band activity (8∼13Hz) in attentional processes (Bae and Luck, 2018; Wolff et al., 2017). This inhibitory role of alpha band activity has been observed in several attention-related domains, including selective attention, working memory, and visual perception (Fukuda and Woodman, 2017; Rihs et al., 2007; Roy et al., 2019). Barne et al. (2020) found that alpha-band power in brain regions contralateral to the expected target location showed greater reduction than in ipsilateral regions. After a target stimulus appears, an increase in alpha-band activity helps to prevent irrelevant memory interference with the current perceptual task (Klimesch et al., 1999). Moreover, lateralization of alpha oscillations is stronger with higher predictive cues, reflecting the involvement and disengagement of functional structures that shape attentional resource allocation (Haegens et al., 2011). In conclusion, alpha-band activity is involved in perceptual processing and shapes the functional structure of the brain to coordinate attentional resources. However, recent studies have reported conflicting findings on the relationship between alpha oscillations and metacognition (such as precision of prediction and confidence). Samaha et al. (2017) recorded trial-by-trial changes in accuracy and discovered a significant negative correlation between alpha power and confidence. In contrast, Benwell et al. (2017) found no relationship between alpha power and perceptual ratings.

In this study, we measured alpha oscillations to assess attentional resource allocation under dynamic environments. We employed a modified Central Cue Posner paradigm where the probability of valid cues fluctuated between three levels (88%, 69%, and 50%), creating a volatile environment. Trialwise belief update trajectories for each participant were estimated by using a hierarchical Bayesian model (Mathys et al., 2011, 2014). Then, we mapped the belief updates to changes in attentional resource allocation using computational modeling (Vossel et al. 2014, 2015), resulting in the attentional resources estimate (α), which was then included as a regressor when analyzing alpha-band oscillations. In this computational model, the amount of attentional resources allocated to the cued location was determined by the precision of the prediction. We examined the relationship between attentional resources (α) and alpha-band oscillations both pre- and post- target across different cue validity conditions. We predicted that attentional resources (α) would predict alpha-band oscillations in the pre-target time phase in a cue-validity dependent manner, with greater prediction when validity was higher. We further predicted that attentional resources (α) would predict alpha-band oscillations in the post-target time period only for validly cued locations.

## Materials and Methods

### Participants

Thirty-one college students participated in this experiment. All participants were right-handed, and had normal or corrected-to-normal vision, and no history of or current neurological impairments. The Ethics Committee of the Institute of Psychology at South China Normal University approved this study. Before the experiment, every participant provided informed consent. Three participants were excluded from further analysis after data acquisition: two participants were excluded due to not following the task instructions, while the other was excluded because of a technical problem with missing marks for the EEG signal. In total, the sample size comprised 28 participants, including 8 males, aged between 20 and 22 years old (mean age: 21.3 years).

### Experiment Procedure

All stimuli were presented on a monitor with a resolution of 1024×768 pixels and a refresh rate of 120Hz, viewed at a distance of 60 cm. All participants were seated in a room shielded from electrical interference. The experimental task was a cue-target paradigm programmed and presented using Psychopy software.

Throughout the course of the experiment, a diamond-shaped fixation point was continuously displayed against a black background (see Figure 1A). On each trial, either the left or the right side of the central diamond-shaped fixation point was highlighted for 200ms as the central cue (2° wide in each visual field, Figure1A). After 400ms, a circular grating (2° wide and 8° eccentric, Figure1A) appeared in a box on either the left or right side of the center as a target stimulus. Participants were required to press the response key immediately after seeing the circular grating. The circular gratings were oriented either vertically or horizontally and were presented for 200ms. Horizontal and vertical Settings are designed to balance physical properties, regardless of reaction. The reaction time window for the participants was set at 1500ms. After the response time window, there was an interval of 800∼1000ms before the next trial commenced. During the experiment, participants were instructed to maintain fixation on the central point and not to respond prematurely.

**Figure 1.**
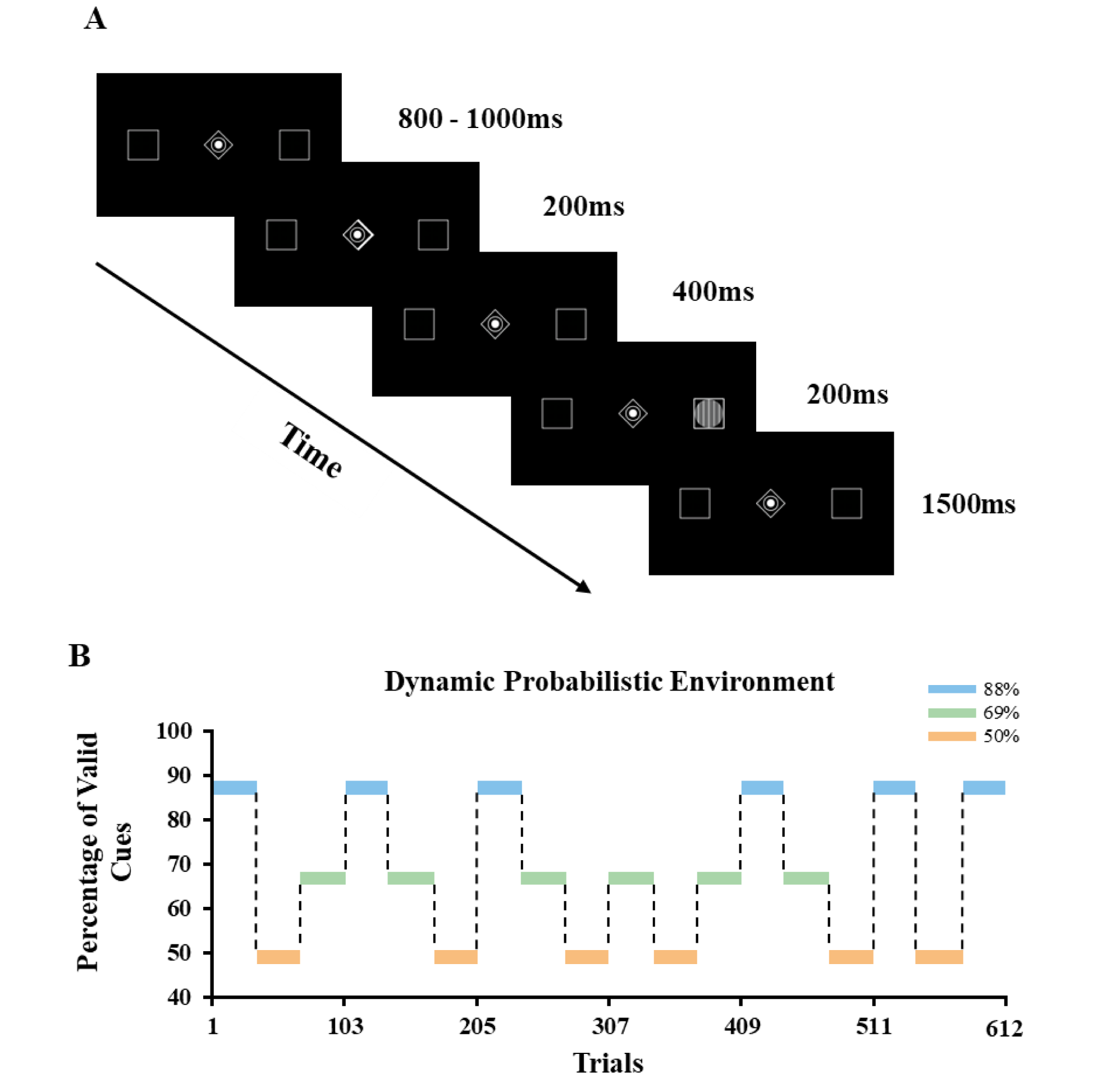
Experimental design. (A) The Central Cue Posner Paradigm was used in this experiment. The valid cue condition shown in the figure indicated that the target was presented at the location pointed to by the arrow. (B) Trial-by-trial changes of cue validities in the manipulated experiment form a fluctuating probability environment.

To ensure that the participants were sufficiently acquainted with and understood the experimental task, they performed 16 practice trials with a constant cue validity of 88% prior to the formal experiment. The formal experiment consisted of 18 blocks, comprising a total of 612 trials, with the participants taking a brief 2-minute break after completing half of the trials. The cue validity, i.e., the probability that the cue accurately indicated the stimulus location, changed unpredictably between 88%, 69% and 50% (see Figure 1B). Each block contained 32 or 36 trials, alternating throughout the experiment. The cue validity within each block remained unchanged. Every block contained an equal number of targets on the left and right side. The total number of blocks featuring each of the three different cue validity levels (88%, 69%, 50%) was the same. The participants were informed only of cue validity changes at the beginning of the experiment, without knowledge of the specific timing or probability of a cue being valid. The participants were all presented with the same trial sequence, which is a standard procedure when studying learning processes involving the inference of conditional probabilities (e.g., Behrens et al., 2007; Dyjas et al., 2012). By keeping the trial sequence constant, we ensure that any variations in model parameters may be ascribed to differences between participants, rather than factors related to the task itself.

### Perceptual and Response Model

In order to capture how individual guides behavior through inference of environmental statistics based on internal perception models, we used a hierarchical Bayesian model established by Mathys et al. (2011, 2014) (open-source code available: https://www.translationalneuromodeling.org/tapas) as the perceptual model (see Figure 2A). We concentrated our analyses on response speed (RS, reponse speed=1/reaction time), because RSs follow a normal distribution. Therefore, trial-by-trial response speed was used to estimate parameters from the perceptual model.

**Figure 2.**
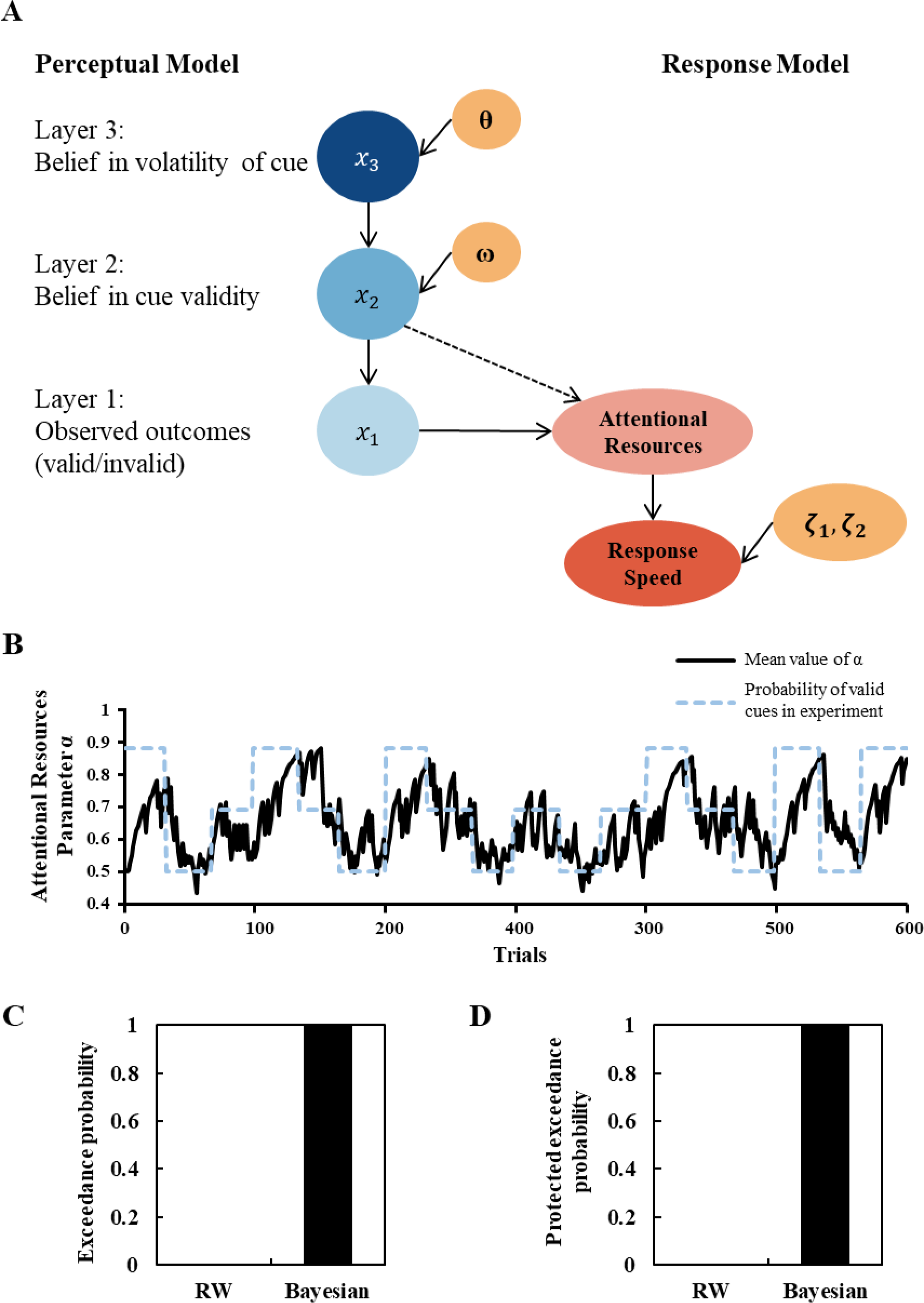
Computational model structure and results. (A) The perceptual model contains three hidden states (x_1_, x_2_, and x_3_) representing the current beliefs of participants. Constant parameters (i.e., ω, θ, 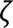_1_ and 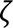_2_) were estimated from each participants’ response speed data. (B) Trial-by-trial changes of the attentional resources parameter α (solid black line) and the true probabilistic environment changes (dashed blue line), with the parameter α calculated on the basis of the mean values over all participants. (C) Bayesian model selection (BMS) results of the tested models: the Rescorla-Wagner model (RW) and the hierarchical Bayesian model. The hierarchical Bayesian model has the best fitting with exceedance probability = 1.0 and (D) protected exceedance probability = 1.0.

In the perception model, x_1_, x_2_, and x_3_ represented the belief updating of three states (for details, seeMathys et al. 2011, 2014). In this experiment, the first layer of the model represented the outcome after the participant observed the target stimulus, indicating whether the cue was valid or not, instead of indicating whether the target stimulus appeared on the left or right side. x_1_ was equal to 1 when cues were valid, and x_1_ was equal to 0 when cues were invalid. The probability distribution of x_1_ values follows a Bernoulli distribution. The second layer of the model, x_2_, represented an individual’s inclination to consider the current cue as valid. When x_2_ was equal to 0, x_1_ had an equal chance of being 1 or 0. As the value of x_2_ approached positive infinity, the probability of x_1_ being equal to 1 approached 1, and the probability of x_1_ being equal to 0 approached 0. The third layer of the model was fit to the logarithmic fluctuations of the effective probability of the cue. The model assumed that the probability distributions of the second and third layers (x_2_ and x_3_) were updated through a Gaussian random walk, where the current trial’s parameter values were determined by the previous trials and the higher levels. Constant parameters, ω and θ, were individual-specific variables. Consistent with prior research (Hein et al., 2021), this study fixed the scaling constant K between the second and third layers to 1.

The perception model depicted the mapping from sensory inputs to subjective probability representations and allowed for obtaining the posterior distribution of the hidden state x at each layer. Prior to observing the x_1_ state, the model used parameters denoted by ∧ to describe an individual’s prediction. Thus, for the current trial t, the posterior belief updating process of the second or third layer was shown in the following equation:

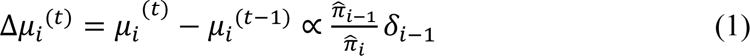

Where 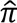_*i*−1_ represented the precision of prediction on the lower-level, 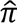_*i*_ represented the posterior belief precision of the current level, and *δ*_*i*−1_ represented the prediction error of the lower level. The derivation process and detailed form of the above formula at specific levels can be found in Mathys et al.’s study (2011, 2014).

The response model described the mapping process from an individual’s posterior belief (generated through fitting the perception model parameters) to the observable response speed. Vossel et al.’s (2014) study compared three models that fit individual response speed and found that the most suitable model was based on the precision of belief in the first level in the perception model, 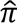_1_^(*t*)^, to calculate an individual’s response speed in the current trial. As shown in Figure 2A, in the response model, the precision 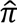_1_^(*t*)^ (= 1/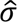_1_^(*t*)^) fit by the first level of the perception model determined the proportion of attentional resources allocated to the cued location, denoted by α, ranging from 0 to 1. In each trial, an individual’s response speed was defined as a linear function of the attention resource parameter α, as shown in the following formula:

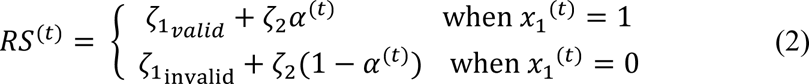

According to the posterior belief probability distribution set by the perception model, the posterior belief probability of the first layer x_1_ followed a Bernoulli distribution. Therefore, when 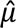_1_^(*t*)^ = 0.5, the value of 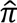_1_^(*t*)^ was equal to 4, which was the minimum value (because 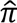_1_^(*t*)^ was the maximum value). To meet the two basic constraints of the response model: (1) the attention resource parameter α should range from 0 to 1, and (2) when 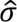_1_^(*t*)^ = 0.5, the attention resource parameter α should also be 0.5 (i.e., when an individual thought that the target stimulus appearing at the cue-directed and uncued locations is equally likely, the attention resources allocated to the cue-directed location should account for half of the total attention resources). At the same time, when the value of 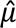_1_^(*t*)^ is less than 0.5, the value of *μ*_2_^(*t*−1)^ will be less than 0. Therefore, under the premise of satisfying all possible values of 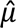_1_^(*t*)^, Vossel et al. (2014) proposed a formula for calculating the attention parameter α:

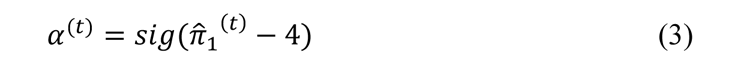

Where the variable 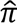_1_^(*t*)^ represented the precision in predicting the upcoming target stimulus at the cue-directed location and was fitted by the first level of the perception model for the current trial t.

The constants ω and θ in the perception model, and the constants 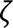_1*valid*_, 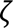_1*invalid*_, and 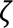_2_ in the response model were fit based on the response time results of each trial for each individual using the variational Bayesian method provided in the Matlab TAPAS toolbox. The constants 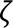_1*valid*_ and 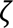_1*invalid*_ represented the individual’s baseline response speed level, while the constant 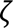_2_ determined the slope of the linear equation, that is, the degree to which an increase in attention resources allocated to the cue-directed location led to an increase in the individual’s response speed.

### Model Comparison

We performed the comparison of model performance between the Rescorla Wagner (RW) model and the hierarchical Bayesian model. We used random-effects Bayesian model selection (BMS) to test which model can better explain the behavioral response. RW model is a simpler learning model based on reinforcement learning theory (Rescorla and Wagner, 1972). It assumes that belief updating is driven by expected errors, but also assumes that the learning rate is fixed. The structure of three-level Bayesian model has been decribed above. It contains not only reinforcement learning theory but also Bayesian learning theory. By comparing the performance of these two models at the group level, it is possible to verify whether participants use updating prediction to adaptively adjust their behavior (Stephan et al., 2009). The BMS indicators to determine the winning model are exceedance probability and protected exceedance probability (codes are open available; Soch and Allefeld, 2018).

### EEG Recording and Time-Frequency Analysis

EEG data was collected using a Neuroscan 64-channel EEG acquisition system, and the arrangement of electrode locations was in line with the 10-20 international standard lead system. The FCz electrode was used as the reference electrode while recording, and the impedance values of all electrodes were reduced to below 5kΩ. During the off-line analysis of EEG data, the average value of the M1 and M2 amplitudes of the left and right mastoid process electrodes was used for re-reference. The sampling rate of EEG was reduced to 500Hz. Low-pass 40Hz digital filtering was performed on the EEG data. EEGLAB toolkit (Delorme & Makeig, 2004) was used to preprocess and analyze EEG data. Independent Component Analysis (ICA) was used to eliminate eye movement artifacts or muscle movement artifacts. Amplitudes beyond ±80μV were marked and automatically eliminated.

Time-frequency analysis was used to analyze the spectral energy of each frequency band of EEG signal. In this experiment, the short-time Fourier transform (STFT) method (Gabor, 1946) was used to process the time-domain signal with a window length of 200ms. EEG signals were segmented from 1000ms before the target to 1500ms after the occurrence of target stimulus. The frequency range included in the analysis was 1 to 30Hz (with 1Hz as a step). Referring to the baseline correction method of Hu et al. (2014), the power value of each frequency band is corrected by subtracting the average power value of the baseline. For the time-frequency analysis before the presentation of the target stimulus, 600∼800ms before the target onset (200ms before the presentation of the cue) was selected as the baseline. For the time-frequency analysis after the presentation of the target stimulus, 300ms before the presentation of the target stimulus was selected as the baseline. For data cleaning, in addition to those trials with abnormal EEG signals, the trials in which the participants made mistakes were also excluded from further analysis.

### General Linear Model Regression Analysis

Regression was used to test the regulatory relationship between model parameters and neural oscillations. Alpha band (8∼13Hz) power values before and after the presentation of the target stimulus within the three cue validity conditions were separately modeled. For the analysis of the time period before the target stimulus, we chose a −600∼0ms locking time window. For the analysis of the time period following the target stimulus, we examined the time interval of 0-500ms after its presentation. Meanwhile, the z-score standardization method was used to process the power values obtained from Fourier transform in the frequency domain. The standardized frequency domain data was then used in the general linear model analysis. The trial-by-trial estimated parameter α from the response model represents the proportion of attentional resources and was used as the regressor across all time points of the electrode of interest.

To test the significance of the regulatory effect of the regression slopes (β) at the group level (i.e., whether the regression coefficient was significantly greater than or less than 0), we performed two-tailed t-tests at each electrode and each time point. In order to exclude false positive results, we conducted an FDR correction (q=0.05) (Benjamini & Hochberg, 1997; Benjamini & Yekutieli, 2001) on the P-value of the regression coefficient (β) for each dimension (time point × electrode point).

## Results

### The Effect of Cue Validity on RS

In order to filter the data included in the experimental analysis, we used the 3sigma principle (Laida criterion) to filter out outliers, which accounted for 0.6% of all data. In addition, incorrect or missing trials were excluded from further analysis, which accounted for 2% of the total number of trials. The analyses were corrected for multiple comparisons using the Bonferroni method.

A 2×3 repeated measures ANOVA, with factors of cue type (valid vs. invalid) and cue validity (88%, 69%, 50%), showed a significant main effect of cue type (*F_(1, 27)_ = 29.1, p < 0.001*), indicating that response speed was faster for targets that were validly cued compared to those that were invalidly cued. The main effect of cue validity was not significant (*p > 0.1*). The interaction effect of cue type×cue validity was also significant (*F_(5, 27)_ = 13.2, p < 0.001*). As shown in Figure 3A, follow-up Bonferroni corrected post hoc comparisons revealed RS differences in different probability environment: The response speed of participants to validly cued targets in the 50% condition was significantly slower than in the 69% and 88% conditions (*p = 0.042* and *p < 0.008,* respectively). Similarly, cue validity probability had a significant effect on response speed for invalidly cued targets, with slower responses in the 88% condition compared to both the 50% and 69% conditions (*p < 0.001* and *p < 0.003,* respectively).

**Figure 3.**
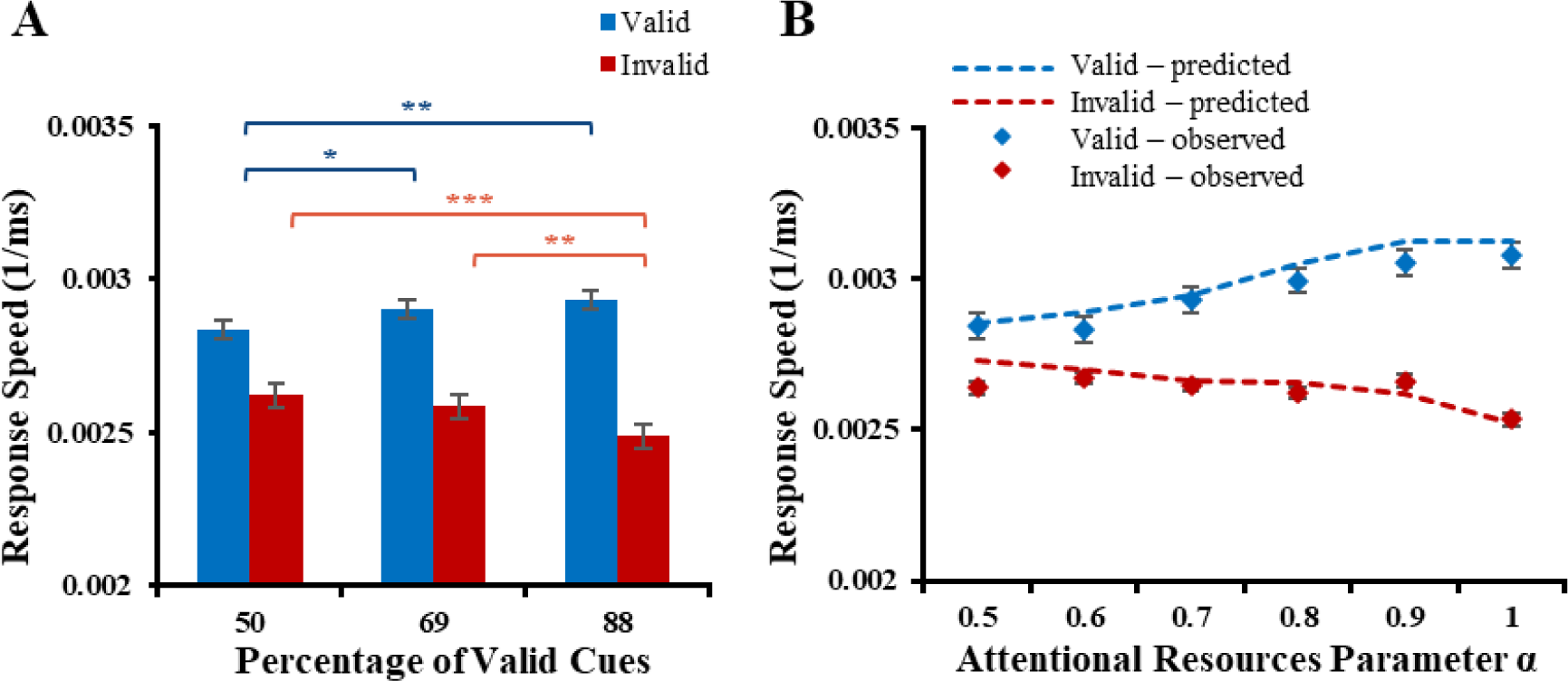
Comparison of (A) response speed of all participants under three distinct probability conditions (response speed =1/ reaction time). Note: *p<0.05, **p<0.01, ***p<0.001. (B) Comparison of predicted and observed response speed relationship with the attentional resources parameter α (i.e., the proportion of attentional resources allocated to the cued location), averaged at the group level.

The response model was used to fit the allocation of attentional resources in each trial, and subsequently to fit the response speed of each participant based on their individual response biases. A repeated measurement ANOVA with a 2×6 design (cue type: valid vs. invalid, binned quantity α: <0.5, .5-.6, .6-.7, .7-.8, .8-.9, .9-1.0) revealed a significant main effect of cue (*F_(1, 27)_ = 112.7, p < 0.001*) and a significant main effect of attentional resources parameter α (*F_(5, 27)_ = 7.2, p < 0.001*). The interaction effect of cue type×attentional resource quantity was also significant (*F_(11, 27)_ = 16.6, p < 0.001*). As depicted in Figure 3B, we compared the predicted reaction speed from the response model with the actual observed reaction speed of the participants. As the attentional resources allocated to the cued location increased, both the predicted and observed reaction speeds exhibited the same pattern of change. This indicates that the combined model can simulate and predict the behavioral performance of the participants realistically. Figure 2B displays the trial-by-trial changes in the attentional resources parameter α throughout the course of the experiment, which were averaged at the group level.

### Model Selection

We used the RW model and the hierarchical Bayesian model respectively to fit the response data of 28 participants. The results of model comparison between these two models were summarized in Figure 2C and 2D. The hierarchical model that earned the highest exceedance probability and protected exceedance probability (with exceedance probability = 1.0, protected exceedance probability = 1.0) was clearly superior.

### Modulation of Precision-Dependent Attention Allocation on Pre-Target Alpha Oscillations Across Three Cue Validities

The analysis of the pre-target time period used regression to characterize the modulation of precision-dependent attentional resources in the alpha band in the three different cue validity probability conditions (88%, 69%, 50%). Cue type (valid versus invalid) was not included in this analysis as there should be no difference in these trial types before the target appears.

The regression results shown in Figure 4A revealed a significant negative relationship between the attentional resources parameter (α) and alpha band power (*P*_*FDR*_*<0.05*) under 88% cue validity. At 138ms prior to the presentation of the target, an examination of the scalp topography shown in Figure 4B reveals the largest effects of attentional resources. The largest effects as calculated by the extreme T-value of the group T-test were concentrated over the right parietal region (with minimum T=-5.43 on P2 electrode). As depicted in Figure 5 (top row), there was no significant relationship observed between the attentional resources parameter (α) and alpha band activity in the 69% and 50% validity conditions. An inspection of the scalp topography (Figure 5, bottom row) also indicated no regional differences for this modulation effect under the conditions of 69% and 50% validity. It should be noted that the statistical analysis included the same number of trials (204 total) for each of the three cue validity conditions and therefore had similar statistical power to detect any differences. These results are consistent with the hypothesis that estimated attentional resources based on Bayesian computational models of the belief updating can account for the deployment of neural attentional resources as measured by the alpha frequency band. Furthermore, this relationship was sensitive to level of certainty, appearing only under conditions of high certainty (88% valid cues).

**Figure 4.**
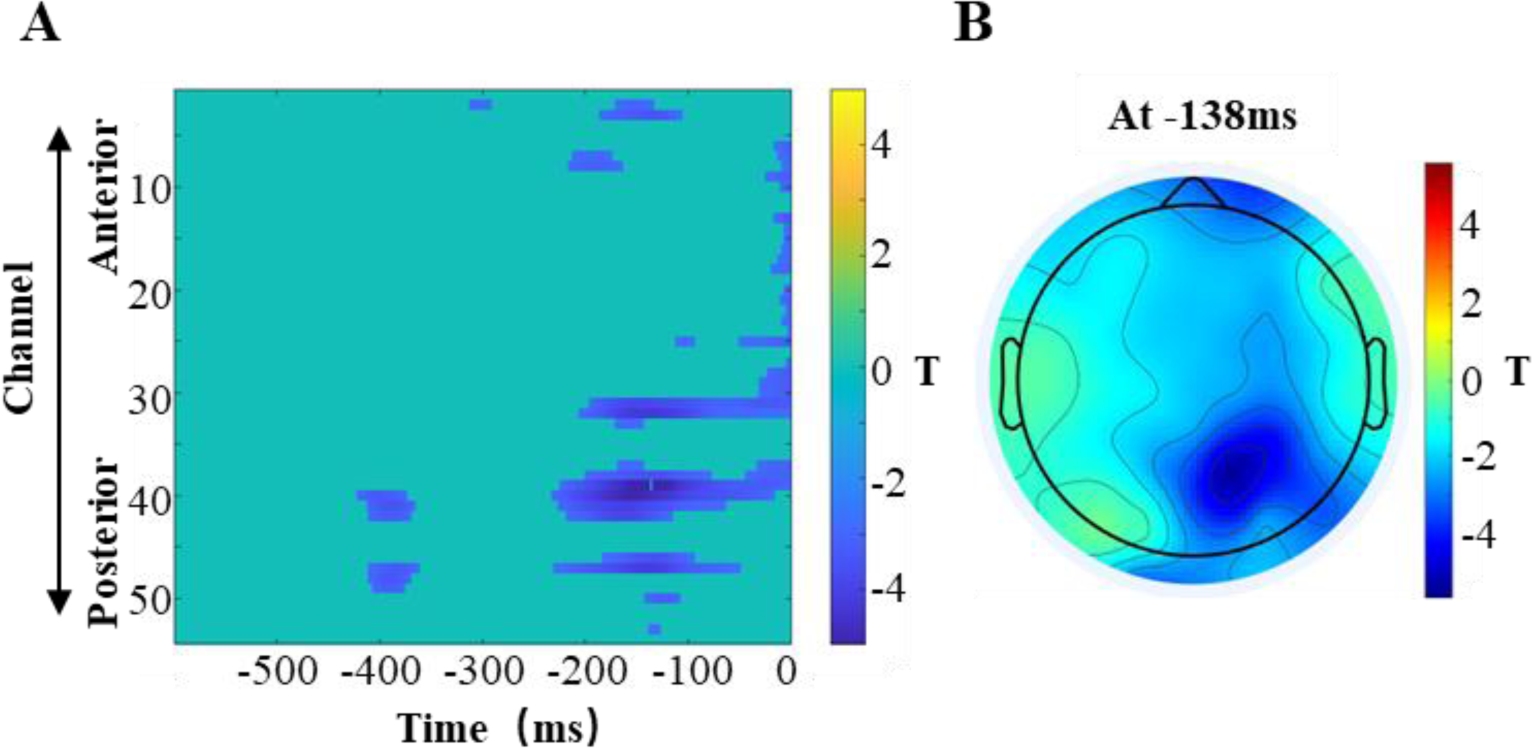
Modulation of pre-target alpha band power in the 88% cue validity condition. (A) The attentional resources parameter (α) significantly and negatively modulated alpha band power in the 600ms time window preceding the presentation of the target (*P*_*FDR*_*<0.05*). (B) At 138ms prior to the presentation of the target, the topographic map presented showed the effect of the attentional resources parameter (α) over parietal electrodes, with the largest effect calculated by the extreme T-value of the group T-test.

**Figure 5.**
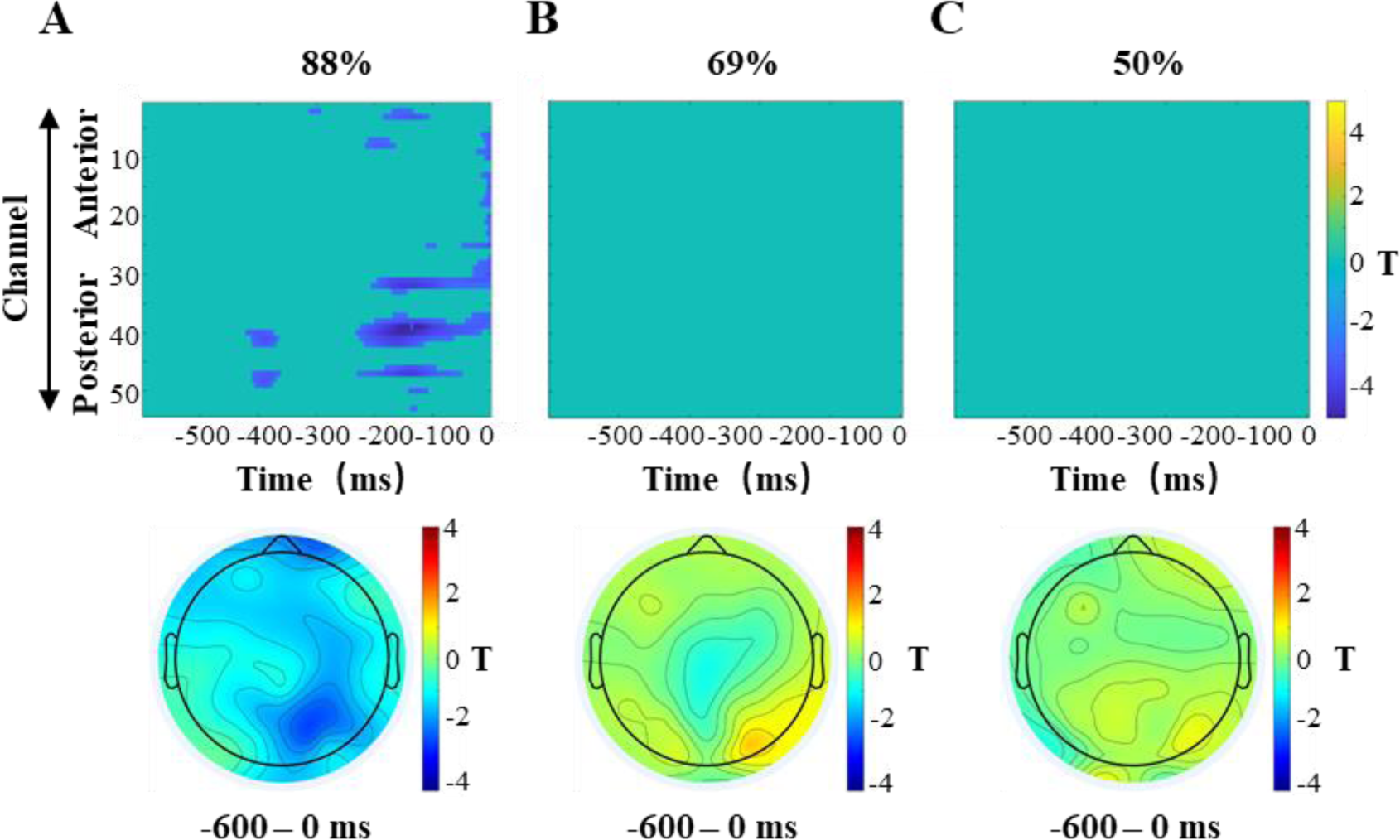
Modulation of pre-target alpha band power across all three cue validities. (A) The attentional resource parameter (α) significantly modulated the pre-target alpha band power under 88% cue validity (*P*_*FDR*_*<0.05*) (first row). Averaging across the entire time span, this modulation was centered over the right parietal region (second row). (B) The attentional resource parameter (α) did not significantly modulate the pre-target alpha band power under 69% cue validity. (C) Attentional resource parameter (α) did not significantly modulate the pre-target alpha band power under 50% cue validity.

### Modulation of Precision-Dependent Attention Allocation on Post-Target Alpha Oscillations between Valid and Invalid Cues

The analysis of the post-target time period also used regression to characterize the modulation of alpha band activity by the attentional resource parameter (α). This analysis focused on differences between valid and invalid cues rather than the three cue validity percentage conditions because there were insufficient trials available for adequate statistical power when the valid and invalid conditions were further subdivided by cue validity. When the target appeared at the location indicated by the cue, the new information provided by the cue in the current trial was useful. In contrast, under conditions of invalid cues, the prediction was erroneous and not useful.

The analysis of the valid cue condition, shown in Figure 6A, revealed a significant negative correlation between the attentional regressor (α) and alpha band power after the appearance of the cued target stimulus (*P*_*FDR*_ *<0.05*). As shown in Figures 6B and 6C, the scalp topography at 40ms and 256ms after the presentation of the target revealed an earlier left central negativity followed by a later right central negativity. The largest negativity was calculated by the extreme T-value of the group T-test (with minimum T=-4.36 at both the C5 and C2 electrodes). Figure 7 compared the valid (A) and invalid (B) cue conditions. In the invalid condition there was no significant modulation observed in the post-target alpha band power by the precision-dependent attentional regressor (α), as shown in the top panel. Furthermore, as shown in the bottom panel, the scalp topography across the entire time period revealed no regional effects within the invalid cue condition.

**Figure 6.**
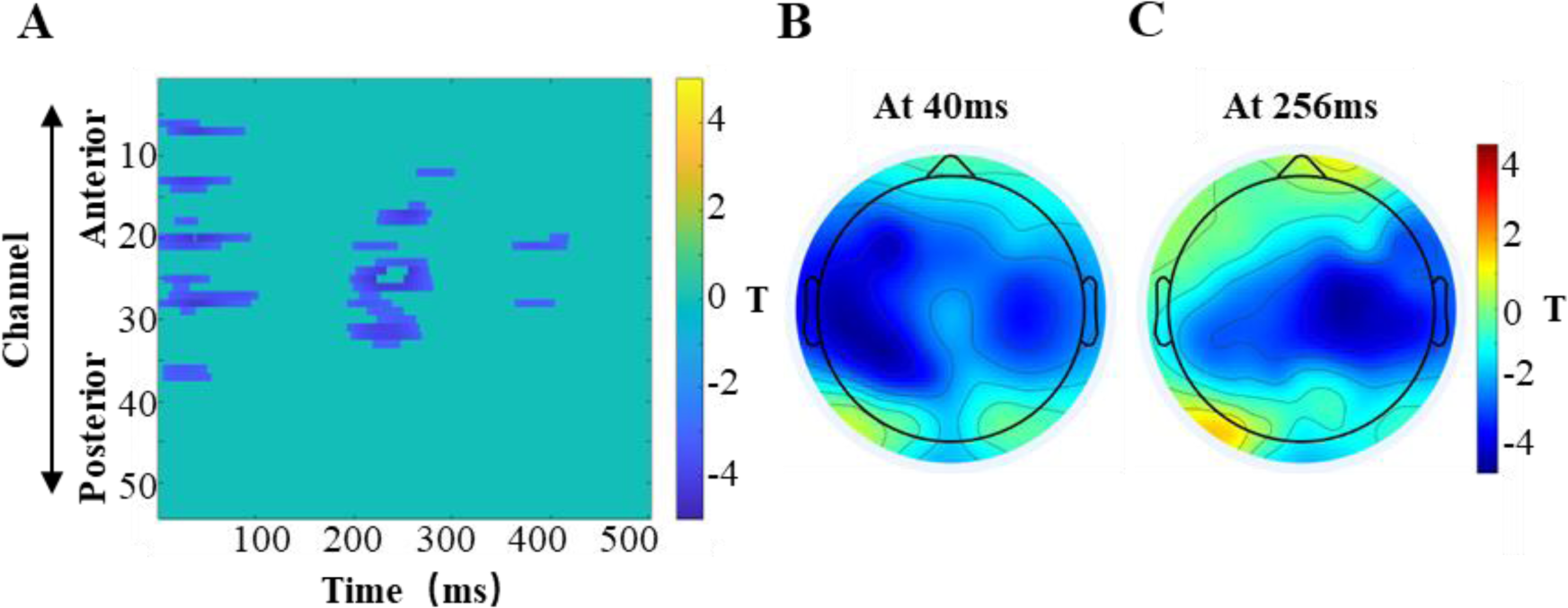
Significant modulation of post-target alpha band power in the valid cue condition. (A) The attentional resource parameter (α) significantly and negatively modulated alpha band power in the 500ms time window after the presentation of the cued target (*P*_*FDR*_*<0.05*). (B, C) At 40ms and 256ms following the target presentation, the topographic maps illustrate the effect of the attentional resource parameter (α) across electrodes, with the most significant effect indicated by the extreme T-value derived from the group T-test.

**Figure 7.**
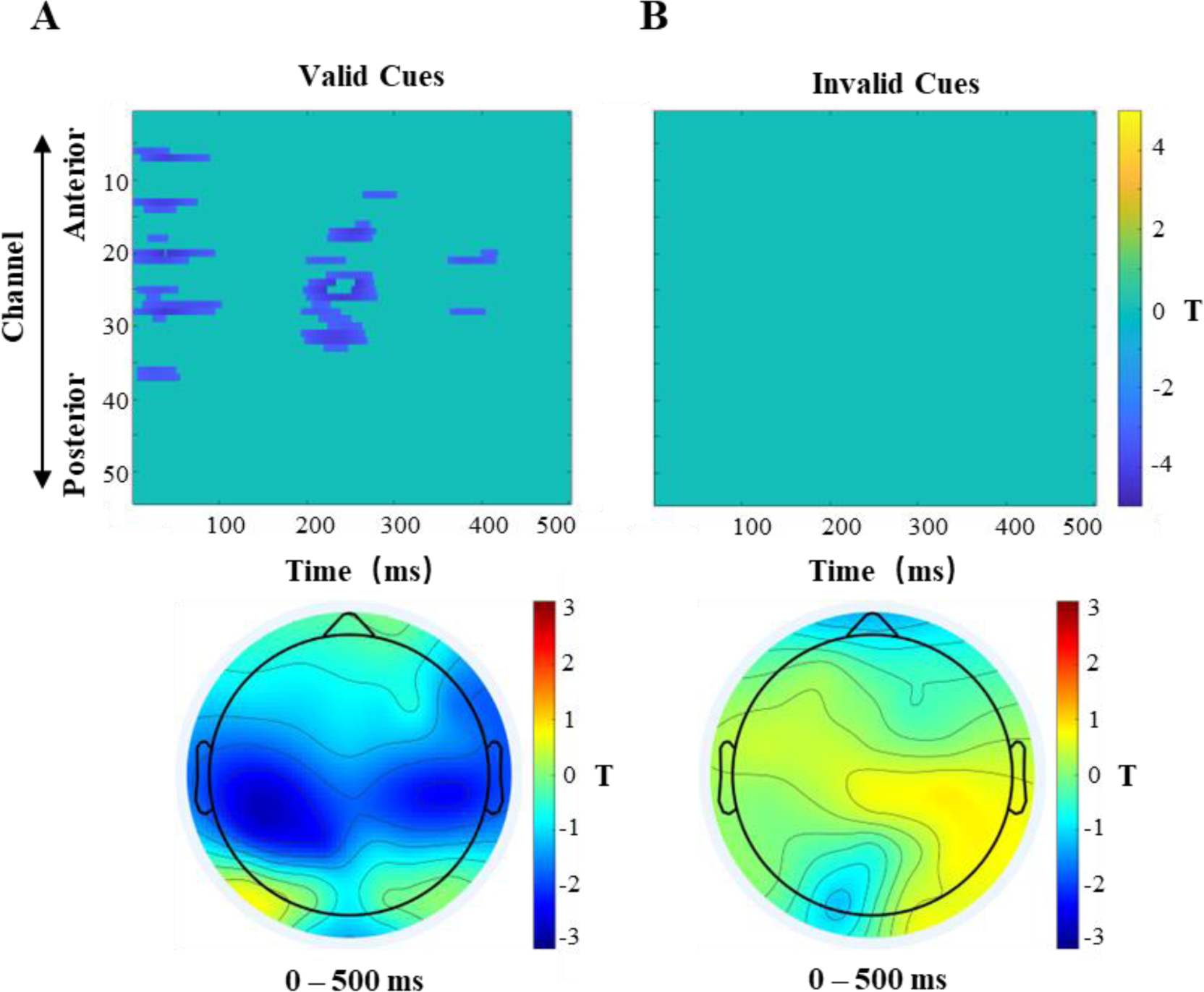
Comparison of the modulation of post-target alpha band power in the valid and invalid cue conditions. (A) Under valid cue conditions, post-target alpha band power was significantly modulated by the attentional resource parameter α (*P*_*FDR*_*<0.05*). (B) Under invalid cue conditions, the attentional resource parameter α did not significantly modulate post-target alpha band power.

## Discussion

This study utilized a combination of Bayesian computational modeling and frequency domain analysis to investigate the computational and neural underpinnings associated with the allocation of attentional resources. The results showed that the attentional resource parameter (α), estimated based on the precision of the predictions, predicted alpha band power related to attention allocation both before and after target presentation. Specifically, before the presentation of the target, the attentional resource parameter (α) negatively predicted alpha band (8-13Hz) power in the high-certainty cue condition (88% valid), whereas this significant relationship was absent in the low-certainty (69% and 50% valid) cue conditions. After the presentation of the target, the attentional resource parameter (α) predicted alpha band power when the stimulus appeared at the cued location (valid cue condition), which manifested as negative correlation with alpha band activity. However, this significant relationship was not found when the prediction was erroneous (invalid cue condition). Overall these findings indicated that attention allocation, which was sensitive to the variation in the probability of valid cues, could account for neural activity in the alpha frequency.

As it was impossible to determine the validity of the cues before the presentation of the target stimulus, we compared the modulatory effects of attentional resources on alpha band activity under three cue validity conditions (88%, 69%, and 50%). As shown in Figure 5, although slight modulatory effects were observed in the 69% and 50% conditions, only in the 88% probability environment did the attentional resource parameter significantly modulate the alpha frequency power. This pattern of results implies that for cues with high certainty (88%), individuals began to deploy attentional resources before the target appeared, which was reflected in alpha band activity.

We used hierarchical Bayesian modelling to characterize the belief updating process. The cognitive parameters derived from the Bayesian model were then used to calculate the proportion of attentional resources in the response model. According to the formula for calculating the attentional resource parameter α, there is a sigmoid relationship between attentional resources and precision of prediction (from the first layer of the perceptual model). As the precision of prediction increases, more attentional resources are allocated to the location indicated by the cue. Therefore, the attention resource measure is precision-weighted or precision-dependent. When the precision of predictions increases, individuals become less sensitive to the incoming evidence and less likely to change their expectations for future trials (Fleming and Daw, 2017; Kiani et al., 2014; Yon et al., 2020). When the surrounding environment is constantly changing, these resulting changes in precision affect the allocation of attention resources, allowing the individual to adapt to the environment. We further found that this adaptation is reflected in alpha band activity.

The present study found that alpha band power before stimulus presentation is sensitive to attentional resources or prediction, which is consistent with previous research findings (Anderson and Ding, 2011; Haegens et al., 2012; Worden et al., 2000). In order to characterize the cognitive functions associated with alpha band power prior to stimulus presentation, many studies using visual detection tasks have reported a significant association between alpha band activity and hit rates (Busch et al., 2009; Mathewson et al., 2009; Romei et al., 2010). However, recent investigations have suggested that this relationship might originate from changes in individual internal standards rather than perceptual sensitivity (Iemi et al., 2017; Limbach and Corballis, 2016). Building upon these findings, Sherman et al. (2016) further explored perceptual sensitivity and discovered that alpha oscillations periodically transmit previous evidence to the visual cortex, thereby modulating the baseline of evidence accumulation. Similar to this finding, Iemi et al. (2017) reported that a decrease in alpha power is associated with a more flexible detection criterion and an increase in baseline excitability, which in turn biases perception toward a higher probability of detecting the stimulus. Our results are in line with several recent studies that found a negative correlation between internal prediction and alpha-band power prior to the appearance of the target stimulus (Baumgarten et al., 2016; Haegens et al., 2012; Samaha et al., 2017; Wöstmann et al., 2019). Our study extended these researches by comparing the results of three cue validity conditions. Only when the cue was highly certain (88% valid probability) did individuals utilize the information provided by the cue to adjust their prediction and inference as measured by the power of neural oscillations in the alpha band. Although attentional resources were still deployed under both low certainty conditions (69% and 50%), the modulation of alpha power could not be explained by changes in attentional deployment in these conditions.

When we examine activity following the presentation of the target stimulus, the presence or absence of the target at the cued location (i.e., differences between valid and invalid cue conditions) becomes a crucial factor. Results revealed that under valid cue conditions (across the 88%, 69%, and 50% probability conditions), the attentional regressor (α) significantly negatively predicted the alpha band power (see Figures 6 and 7). However, under invalid cue conditions there was no significant correlation between the parameter and the power of alpha band frequency (see Figure 7). These results suggest that when the target stimulus appeared at the cued location, participants utilized prior predictions to modulate alpha-band activity to facilitate task performance. Consequently, when the target stimulus failed to appear at the cued location, the lack of a significant correlation between prior predictions and alpha-band activity may suggest the need for a recalibration of attentional resources (Schröger et al., 2015; Wang and Shen, 2018).

Early research on alpha frequency activity found that when participants were in a resting state before the appearance of stimuli (i.e., not in a task state), neural activity in the alpha power increased, whereas after the onset of sensory stimuli, neural activity in the alpha frequency decreased. Therefore, researchers believe that synchronized oscillatory neural activity in alpha oscillations is associated with negatively activated cortical areas (Pfurtscheller et al., 1996). Recently, an increasing number of electrophysiological studies have updated our understanding of the functional role of neural activity in alpha oscillations, suggesting that alpha oscillations play a positive modulatory role in cognitive processing (Haegens et al., 2014; Jensen and Mazaheri, 2010; Klimesch, 2012). Some researchers have associated neural activity in alpha oscillations with inhibitory actions, suggesting that alpha oscillations suppress sensory states unrelated to the task (Doesburg et al., 2009; Klimesch et al., 1999; Stern, 2002). To successfully execute a task, not only do cognitive processes related to the task need to be facilitated, but the cognitive control system also needs to inhibit the cognitive activities that interfere with task performance. This inhibitory function is thought to be achieved through alpha oscillations generated in cortical areas unrelated to the task. For instance, a magnetoencephalography (MEG) study conducted by Jokisch and Jensen (2007) found that during face recognition tasks, alpha oscillations decreased in the ventral visual pathway, but increased in the dorsal visual pathway. Another MEG study by Hwang et al. (2014) revealed that the increase in alpha power in the frontal eye field (FEF) inhibited stimulus-driven attention that was unrelated to the task. The conclusions of these studies can effectively explain the negative correlation between attentional resources and alpha oscillations found in our results. When the target stimuli appeared at the cued location, an increase in alpha oscillation power may reflect the suppression of irrelevant information by cognitive processing if the individual has fewer allocated attentional resources at that time (which could potentially indicate distraction or other factors).

Our results extend previous attentional cueing researches from two perspectives. First, we verified the impact of the precision of prediction on attentional resource allocation (Vossel et al. 2006, 2014, 2015; Feldman and Friston 2010). Secondly, we further compared valid and invalid cue conditions. Consistent with previous research, a large body of literature has found differences in cognitive processes such as attentional resource allocation and interference suppression between valid and invalid cue conditions (Klimesch, 2012; Pützer et al., 2019; Sawaki et al., 2012). For instance, Bonato et al. (2016) found that when cues were predictive, the reaction time difference between valid and invalid trials was significantly increased. Based on our findings, a valid cue results in more informative predictions and elicits negative modulation of alpha oscillations. Conversely, an invalid cue leads attentional resource allocation to become more dispersed, and modulation of alpha oscillations may fail to occur.

Recent studies have proposed that alpha oscillations can regulate the transmission and processing of neural activity, and thus contribute to neural activity in regions including the visual cortex, motor cortex, prefrontal cortex, temporal lobe, and parietal lobe (Huang et al., 2021; Labree et al., 2020; Luft et al., 2018; Wianda and Ross, 2019). Based on the scalp topographical maps in our studies, it was evident that these modulatory effects were distributed across different cortical regions before and after target presentation. Specifically, the modulatory effect before target presentation was present on the scalp above the parietal cortex, predominantly on the right hemisphere. In contrast, the modulatory effect after target presentation was distributed in acriss bilateral central scalp locations, with a leftward predominance in the first 100ms and a rightward predominance after 200ms. Zhou et al. (2021) found that prestimulus alpha band power in the occipital region could modulate behavioral accuracy. Vossel et al. (2015) found that attention following target presentation modulated connectivity strength between the right temporo-parietal junction (rTPJ) and the right frontal eye field (FEF), as well as connectivity between the rTPJ and the putamen, reflecting deployment of attentional resources via brain network connectivity. Our results are consistent with these results. These studies collectively suggest that the brain regions involved in attention allocation based on Bayesian belief updating are different before and after target presentation, reflecting the information transfer among brain regions. Future studies in this area may benefit from using magnetoencephalography (MEG) and related techniques, which combine greater spatial resolution with high temporal precision, to investigate the relationship between alpha oscillations and flexible attention allocation.

In conclusion, this study reveals that Bayesian belief updating for the allocation of attentional resources is reflected in alpha band power. The precision of prediction affects the allocation of attention resources. In a dynamic environment, the modulatory relationship between precision-dependent attentional resources and alpha band activity is sensitive to cue validity. These results contribute to our understanding of how attention resources are allocated adaptively and reflected in alpha oscillations.

